# Transcriptional kinetics of X-chromosome upregulation

**DOI:** 10.1101/540104

**Authors:** Anton J. M. Larsson, Christos Coucoravas, Rickard Sandberg, Björn Reinius

## Abstract

Ohno’s hypothesis postulates that X-chromosome upregulation rectifies X-dose imbalance relative to autosomal genes, present in two active copies per cell. Here we dissected X-upregulation into kinetics of transcription, inferred from allele-specific single-cell RNA-sequencing (scRNAseq) data from somatic mouse cells. We confirmed increased X-chromosome expression, and remarkably found that the X-chromosome achieved upregulation by elevated burst frequencies. This provides mechanistic insights into X-chromosome upregulation.

---

In therian mammals, the X-chromosome is present as two copies in female cells while one copy in male cells. The X and Y-chromosomes evolved from an ancestral autosomal pair on which the genetic sex-determining factor *Sry* appeared around 166 million years ago^1^. To preserve genetic linkage between *Sry* and Y-linked male-beneficial genes, Y-X recombination was repressed, leading to Y-chromosome degeneration and gene loss over time^2^. This rendered males monosomic for X-linked gene products and thus imbalanced with the diploid autosomal (A) part of the genome. Landmark theoretical work^3^ by Susumo Ohno proposed that cells restore X:AA balance by doubling the expression of chromosome X, resulting in the X:AA expression ratio of 1 (analogous to X:A ratio 2). This concept (**Figure 1a**) is termed Ohno’s hypothesis, and the emergence of this compensatory process is presumed to predate the evolution of X-chromosome inactivation^4^ (XCI) by which female cells silence one X-chromosome to reach male-equivalent expression levels for X-linked genes.

**Figure 1.**
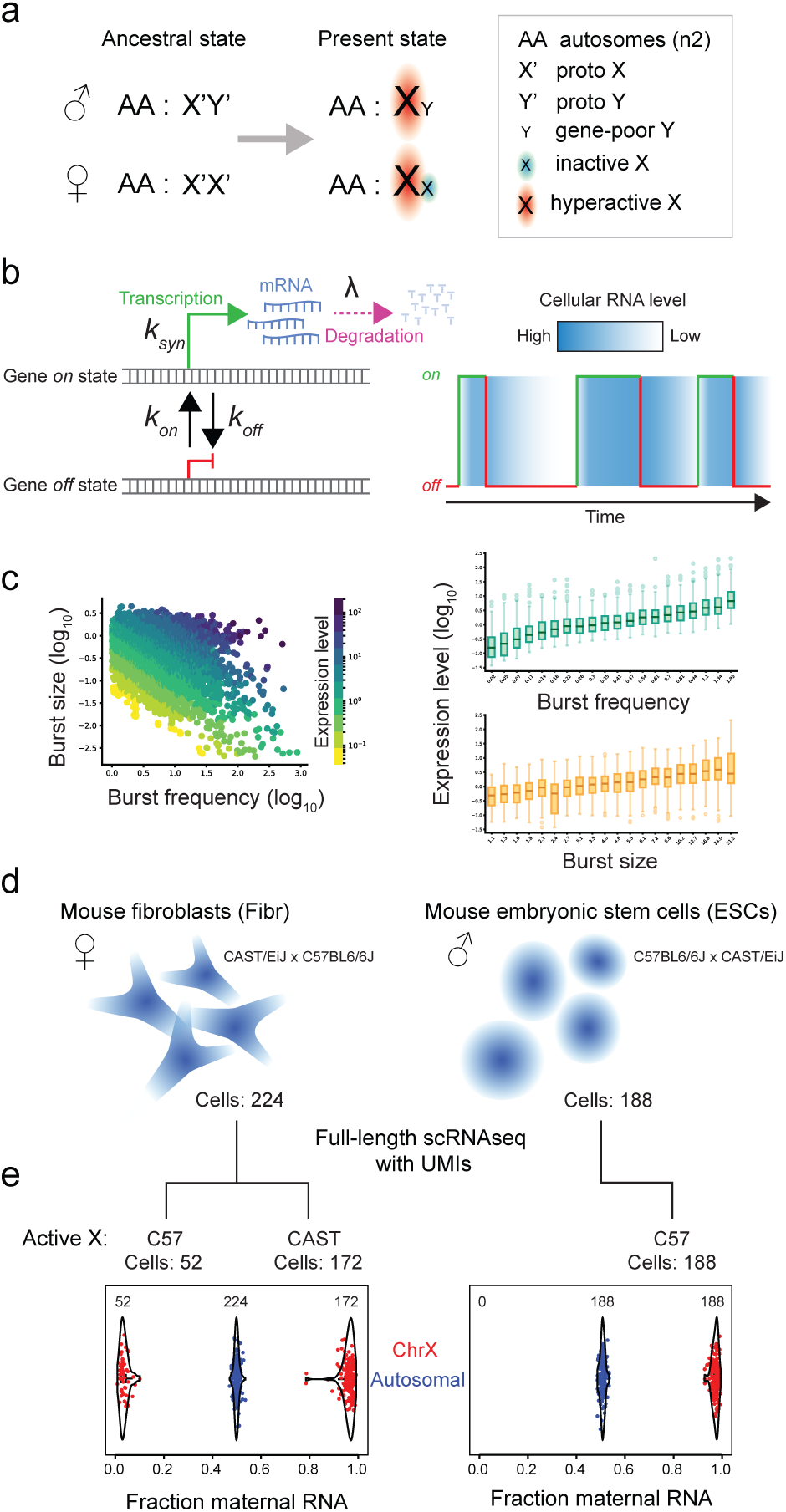
Investigating X-chromosome upregulation by transcriptional kinetics. **(a)** The concept of Ohno’s hypothesis: Relative levels of gene dose between autosomal and X-linked genes may have been restored by chromosome-wide upregulation of X-linked genes. **(b)** Leti: The stochastic process used to model transcriptional bursting in this study. Right: Transcriptional bursting results in fluctuations in levels of cellular RNA. The time the gene is spent on and off are colored in green and red respectively **(c)** Leti: Burst frequency (*k*_*on*_) and burst size (*k*_*syn*_/*k*_*off*_) inferred in genes on the C57 allele in cells expressing the C57 X-chromosome in primary mouse fibroblasts, colored by expression level (n=6579 genes). Right: Burst frequency (top) and size (botiom) both determine the expression level of genes (20 bins, 329 genes per bin). **(d)** The type, sex, and genotype of the mouse cells analyzed in this study. **(e)** Segregation of cells based on expressing either the maternal or paternal X-chromosome due to XCI in female cells. Each dot represents a cell and the fraction RNA molecules originating from the maternal allele is shown for chromosome X (red) and autosomes (blue).

Several studies have reported X-upregulation in mammals, initially using microarrays^5,6^ and later using RNA-sequencing^7,8^. But the validity of Ohno’s hypothesis has also been contested^9-11^, and a caveat to bulk analysis is that cellular heterogeneity may skew estimates due to cell subpopulations expressing chromosomes unequally^7^. Emerging scRNAseq technology provides the opportunity to asses X:A dose balance at the level of the actual regulatory system (i.e. the cell); but dedicated scRNAseq studies are so far few^11-14^, and some arrived at opposite conclusions regarding X-upregulation from the very same scRNAseq data^11,12^. Moreover, a fundamental limitation in all present scRNAseq analyses is that they lack unique molecular identifiers (UMIs) needed to avoid library amplification bias (which is particularly severe for scRNAseq cross-gene comparisons)^15^, or, lack the full-length read coverage of transcripts needed for allele-specific and single-gene-copy analysis. Finally, eukaryotic gene expression occurs through episodic bursting of RNA synthesis^16-19^, but kinetic studies of X-upregulation are completely lacking. Here, we dissected X-upregulation using transcriptional kinetics, providing mechanistic insight to this process.

We investigated primary female fibroblasts (n=224) and male embryonic stem cells (ESCs, n=188) from outbred mice (hybrids of CAST/EiJ and C57BL/6J) that were subjected to full-length scRNAseq with UMIs^18^ (**Online Methods**). We recently developed methods to infer transcriptional burst parameters individually for each gene copy^18^ using the two-state model of transcription^20^, providing allele-specific estimates of burst frequency (*k*_*on*_) and burst size (*k*_*syn*_/*k*_*off*_) (**Figure 1b-c**). In order to explore the kinetics of X-upregulation, we first segregated female fibroblasts expressing either the maternal (CAST, n=172) or paternal (C57, n=52) X-chromosome due to random XCI. Expectedly, male ESCs (n=188) transcribed exclusively the maternal X-allele (**Figure 1d-e**). To investigate X-chromosome expression per cell, we first counted mRNAs *de facto* present in the individual cells (i.e. ≥1 UMI) and calculated cellular expression levels for genes on the active X and autosomal chromosomes. Strikingly, cellular X-chromosome RNA levels were higher than those of autosomes within the same cells (*P*≤ 1.2×10^-9^, paired Wilcoxon rank-sum test) (**Figure 2a**). To avoid possible confoundment of few high-expressed genes dominating chromosomal expression estimates we excluded genes beyond the 95^th^ percentile of expression levels per cell. Other thresholds such as the 90^th^ and 99^th^ percentile provided similar results (**Figure 2b**) and the X:A difference was significant across a wide span of thresholds (**Figure 2c**). Next, we determined gene-wise expression-level distributions for chromosome X (n=149 genes, fibroblast C57 allele) and autosomes (n=3790 genes, fibroblast C57 allele), including only expressed genes (average ≥1 UMI detected per cell) that also had robustly inferable kinetic parameters (**Online Methods**). This revealed a positively shifted expression-level distribution for X-linked genes (median fold-change: 1.4, *P*= 4,2×10^-5^, two-sided Wilcoxon test; fibroblast C57 allele) (**Figure 2d-e**). Comparing distributions for all chromosomes, we observed such a shift to be unique for chromosome X (**Figure 2f-g**). These results were replicated over the fibroblast CAST allele and male ESCs (**Supplementary Figure 1-2**). This confirms Ohno’s hypothesis.

**Figure 2.**
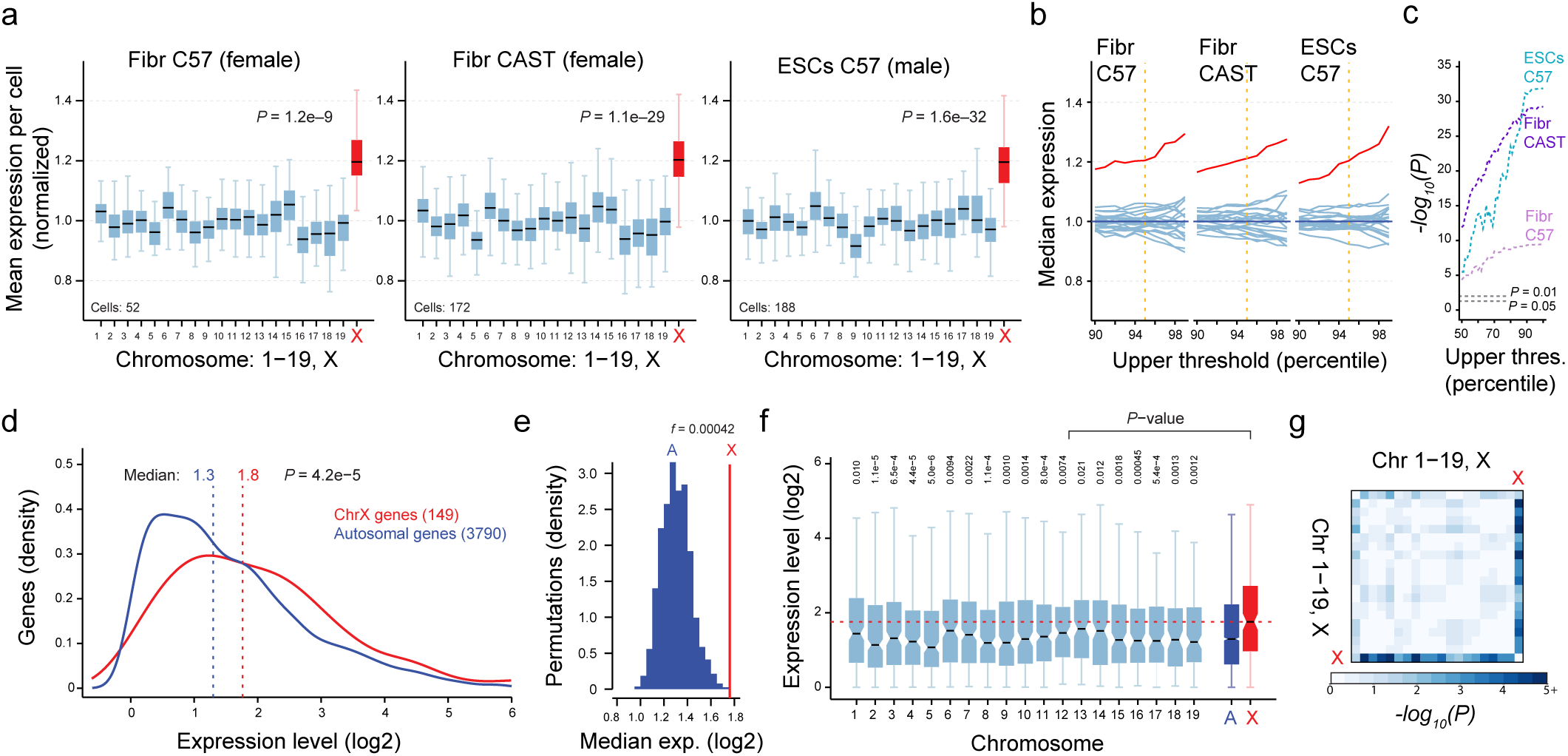
Expression levels of X-linked genes are higher than those of autosomal genes. **(a)** Cellular mean expression of genes on each chromosome normalized by mean expression of all autosomal genes in each cell and allele shown as boxplots. Genes beyond the 95th percentile in expression level removed. Centre lines denote the median; hinges denote the first and third quartiles; whiskers denote 1.5× the interquartile range (IQR). Wilcoxon rank-sum test was used for significance testing. **(b)** Median of normalized cellular expression levels for chromosome X (red) and autosomes (light blue), and **(c)** *P*-values; shown for different upper thresholds. **(d)** Distribution of expression levels of autosomal and X-linked genes on the C57 allele in fibroblasts. Wilcoxon rank-sum test was used for significance testing. **(e)** Histogram of median expression levels of randomly subsampled autosomal genes compared to the median of X-linked genes (n=149 genes; 100,000 permutations), and *f* denoting fraction permutations for which the autosomal median reached that of chromosome X. **(f)** Distribution of expression levels for genes on each autosomal chromosome (light blue), all autosomes (dark blue), and chromosome X (red). Centre lines denote the median; hinges denote the first and third quartiles; whiskers denote 1.5×IQR. One-sided Wilcoxon rank-sum test was used for significance testing. **(g)** Heatmap of *P*-values from all pairwise comparisons of expression levels between chromosomes using two-sided Wilcoxon rank sum test.

In the two-state model of transcription, expression levels are determined by *(1)* the fraction of time the gene spends in transcriptional on-state, generating a burst of RNA copies; *(2)* the average number of RNA molecules synthesised during such a burst; and *(3)* the degradation rate of RNA (**Figure 1b-c**). We compared burst frequencies (*k*_*on*_) for chromosome X and autosomes. Intriguingly, X-chromosome genes maintained distinctly elevated burst frequencies (median fold-change: 1.5, *P*= 3.3×10^-7^, Wilcoxon test, fibroblast C57 allele, **Figure 3a-d**). This was validated over the fibroblasts CAST allele as well as in male ESCs (*P*= 3.7×10^-6^ and 1.1x 10^-7^, respectively) (**Supplementary Figure 1-2**). Next, we performed similar analyses for burst sizes (*k*_syn_/*k*_off_) and observed the lack of significant difference between chromosome X and autosomes (*P*= 0.93; power: 99%, **Figure 3e-h** and **Supplementary Figure 1-3**). To confirm that these kinetic features were not unique for fibroblast- or ESC-specific transcripts, we repeated the analyses using housekeeping genes (**Online Methods**) (**Supplementary Figure 4**). Altogether, our results imply that X-chromosome upregulation occurs through increased frequency of transcriptional bursting in both female and male cells.

**Figure 3.**
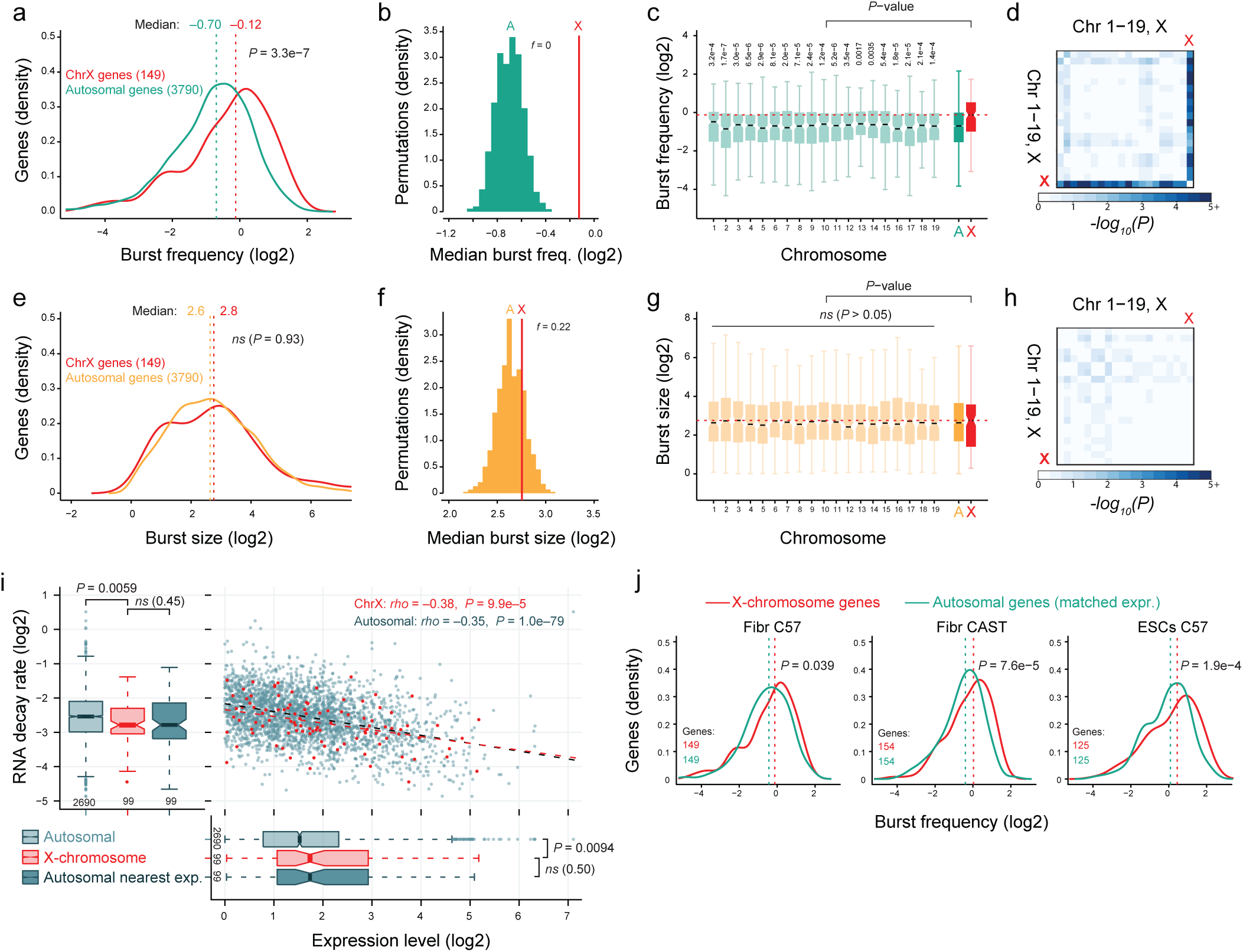
Burst frequencies of X-linked genes are higher than those of autosomal genes but not burst sizes. **(a)** Distribution of burst frequency (*k*_*on*_) of autosomal and X-linked genes on the C57 allele in fibroblasts. Wilcoxon rank-sum test was used for significance testing. **(b)** Median burst frequency of randomly selected subsets of autosomal genes compared to the X-linked genes (n=149 genes; 100,000 permutations), and *f* denoting fraction permutations for which the autosomal median reached that of chromosome X. **(c)** Distribution of burst frequency for each autosomal chromosome (light green), all autosomal genes (dark green), and chromosome X (red). Centre lines denote the median; hinges denote the first and third quartiles; whiskers denote 1.5×IQR. One-sided Wilcoxon rank-sum test was used for significance testing. **(d)** *P*-values from all pairwise comparisons of burst frequency between chromosomes using the Wilcoxon rank sum test. **(e-h)** Same as (a-d) but for burst size (*k*_*syn*_/*k*_*off*_). **(i)** Top leti: Distributions of RNA decay rate for autosomal, X-linked and expression-matched autosomal genes in the C57 allele in ESCs. Top right: Correlations between RNA decay rate and mean expression level (n=2690 autosomal and 99 X-linked genes, spearman correlation). Botiom: Distributions of expression levels for autosomal, X-linked and expression-matched autosomal genes. Centre lines denote the median; hinges denote the first and third quartiles; whiskers denote 1.5×IQR. One-sided Wilcoxon rank-sum test used for significance testing. **(j)** Distribution of burst frequency for X-chromosome (red) and autosomal genes of matched (nearest) expression value, shown for each allele in fibroblasts and the C57 allele for ESCs. Paired one-sided Wilcoxon rank-sum test was used for significance testing between expression matched autosomal and X-genes.

Previous studies^21,22^ found that X-chromosome transcripts tended to have longer RNA half-lives than those of autosomes, suggesting that lowered decay rates contribute to dosage compensation. We used metabolic-RNA-labelling data from mouse ESCs (**Online Methods**), and investigated RNA decay rates for X-linked and autosomal genes. This indeed confirmed lowered decay rates for X-chromosome transcripts (*P*≤ 1.3×10^-4^, **Figure 3i**). However, we also observed the general trend that expression levels negatively correlated with decay rates independently of chromosomal origin of RNAs (Spearman correlation –0.38 and –0.35; *P*= 9.9×10^-5^ and *P*= 1.0×10^-79^ for X and autosomal genes, respectively; **Figure 3i**). We tested whether X-genes had lowered decay rates given their expression levels by calculating the distribution of decay rates for autosomal genes of same expression levels. X-chromosome genes and matched autosomal genes had similar decay rates (medians 0.15 and 0.15, *P*= 0.50, paired Wilcoxon test) (**Figure 3i**). This does not conflict that increased RNA stability might contribute to dosage compensation, but does motivate the search for X-specific regulatory features. We reasoned that if increased burst frequency is a key mechanism to achieve X-upregulation, a shifted burst-frequency distribution might be detectable even when compared to autosomal genes of matched expression levels, albeit expectedly with a smaller shift since the autosomal expression distribution would artificially be shifted to higher values. Indeed, this was valid for both fibroblast alleles as well as for ESCs (*P*= 0.039; 7.6×10^-5^; 1.9×10^-4^, respectively; paired one-sided Wilcoxon test) (**Figure 4j-k**).

Here we investigated X-chromosome upregulation at the resolution of transcriptional kinetics for the first time. Using scRNA-seq data with both full-transcript coverage and UMIs, we simultaneously obtained precise estimates of expression levels and detailed allele-specific information for single-gene-copy inferences. Analysing these novel data, we confirmed the validity of Ohno’s hypothesis by detecting positively shifted expression-level distributions for chromosome X both at cellular and gene-wise level (**Figure 2** and **Supplementary Figure 1a-b**). The observed ∼1.4–fold change of X:A levels (**Figure 2c** and **Supplementary Figure 1-2**) rather than the theoretical 2-fold change suggests that not all X-linked genes attained dosage compensation, or that less than 2-fold upregulation is adequate to achieve sufficient X:A balance for most genes.

While XCI has been extensively explored, providing profound insights into gene silencing, mechanistic correlates to mammalian X-upregulation are few^13,21,22^. By breaking down expression levels into kinetic parameters of transcription, we showed for the first time that dosage compensation is achieved by increased burst frequencies on chromosome X. Since burst frequencies are preferentially encoded in enhancer elements^18^, it is likely that the increased transcriptional output from the X-chromosome is due to trans-acting factors affecting enhancers. Moreover, such factors would preferentially target enhancers located on the X-chromosome.

## Methods

### Sequencing and classification of expressed X-chromosome allele in each cell

A modified version of Smartseq2^23,24^ was used to sequence the transcriptomes of primary mouse tail fibroblasts (n=224) and ESCs (n=188), as described in Larsson et. al.^18^. In brief, we utilized a modified Smartseq2 strand-switch primer containing a UMI (6-base random sequence). Sequencing was performed on an Illumina Nextseq 500, and sequence reads were mapped to the C57 and CAST genomes, and reads spanning strain-specific SNPs were counted as described previously^18,25,26^. The sequencing data is available at <u>E-</u> <u>MTAB-7098</u>. In order to classify cells as either having the C57 or the CAST X-chromosome active in the female fibroblasts, due to X-inactivation, we calculated the aggregated expression of all genes and alleles on the X-chromosome and compared the sums. The parental X-chromosome with the higher sum was classified as having that allele active. In addition, we calculated the fraction of maternal RNA as shown in **Figure 1e** to further verify this classification. All male ESCs expressed only the maternal X chromosome, as these cells carry only one X-chromosome copy.

### Inference of transcriptional kinetics parameters

In the model of transcription used, a gene switches to the on state and off state at exponentially distributed times with rates *k*_*on*_ and *k*_*off*_ respectively. When the gene is on, RNA is produced at rate *k*_*syn*_. Regardless of gene state, the RNA is degraded with rate *λ*. In brief, the probability distribution for the steady state of the stochastic process illustrated in **Figure 1b** is the Beta-Poisson compound probability distribution. The parameters of this process were estimated in the time scale of degradation (*λ*) by maximum likelihood. Detailed information regarding the inference procedure is described in Larsson et. al.^18^. For cells in which the C57 X-chromosome allele was active, we inferred the parameter for autosomal genes at the C57 allele in the same cells (n fibroblasts C57= 52, n ESCs C57= 188). For cells in which the CAST X-chromosome allele was active, we inferred the parameter for autosomal genes at the CAST allele in the same cells (n cells fibroblast CAST= 172). RNA decay rates were obtained for mouse ECSs from Supplementary Table 1 of Herzog et. al.^27^.

### Comparisons between chromosomes

In all comparisons, we included genes that were expressed (≥1 UMI) in the given cells and that also had robustly inferable kinetic parameters (defined as: within the bounds of the maximum likelihood procedure (10^-3^ < *k*_*on*_, *k*_*off*_ < 10^3^ and 1 < *k*_*syn*_ < 10^4^) and biologically feasible 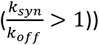). For the cell-wise analysis of chromosome expression levels (**Figure 2a**) we included genes that were detected (at least 1 UMI) in each individual cell and excluded genes beyond the 95^th^ percentile of expression levels per cell. We calculated the mean expression level for each chromosome in each individual cell and divided these values by the mean expression of all autosomal genes in the individual cells (resulting in normalized cellular ChrN/ChrA values). We repeated the analysis using different upper expression thresholds, and cellular X:A differences were significant even when excluding as much as the upper 50% of genes (**Figure 2c**). To compare gene-wise expression levels between chromosome X and autosomes, we calculated the mean expression for each gene across the cells of given cell-type and genotype (C57 allele in cells having the C57-X-chromosome active, and CAST allele in cells having the CAST-X-chromosome active) and plotted a smoother kernel density. To assess the probability of drawing a set of genes with a median equal to or higher than that of X-linked genes from the autosomal distribution, we randomly subsampled autosomal genes (n= n_[X-linked genes]_; e.g. 149 in case of fibroblast C57 allele) and calculated the median expression level. This was repeated 100,000 times (**Figure 2e**), and we calculated the fraction (*f*) permutations for which the median of the random autosomal subset equated or exceeded the median of chromosome X. Corresponding analyses were performed for burst frequency (*k*_*on*_), and burst size (*k*_syn_/*k*_off_). To assess the differences between chromosomes in mean expression (UMIs per cell), burst frequency, and burst size, we used the Wilcoxon rank-sum test as specified in the figure legends for each comparison. Housekeeping genes used in **Supplementary figure 4** were ubiquitously expressed genes^28^ (genes expressed across 17 mouse tissues).

### Power analysis of detecting X-chromosome wide upregulation in burst size

To assess whether we would have the power to detect a 1.4x change in expression level due to burst size, we randomly selected 149 autosomal genes, increased their burst size by 1.4, used the Wilcoxon rank-sum test and noted the *P*-value of this test. This was repeated 1000 times.

### Code availability

The computational code used for calculations and plotting of data are available at https://sourceforge.net/projects/ kinetics-of-x-upregulation.

## Acknowledgements

This study was supported by grants from the Ragnar Söderberg Foundation, the Swedish Research Council (2017- 01723), and Åke Wiberg’s Foundation to BR.

## Author contributions

BR conceived the study. AL and BR analysed the data and wrote the manuscript. AL, CC, RS and BR participated in interpreting the data and editing the manuscript.

## Online content

Supplementary figures 1-4.

