## Supplementary figures for "Transcriptional kinetics of X-chromosome upregulation"

Supplementary figure 1

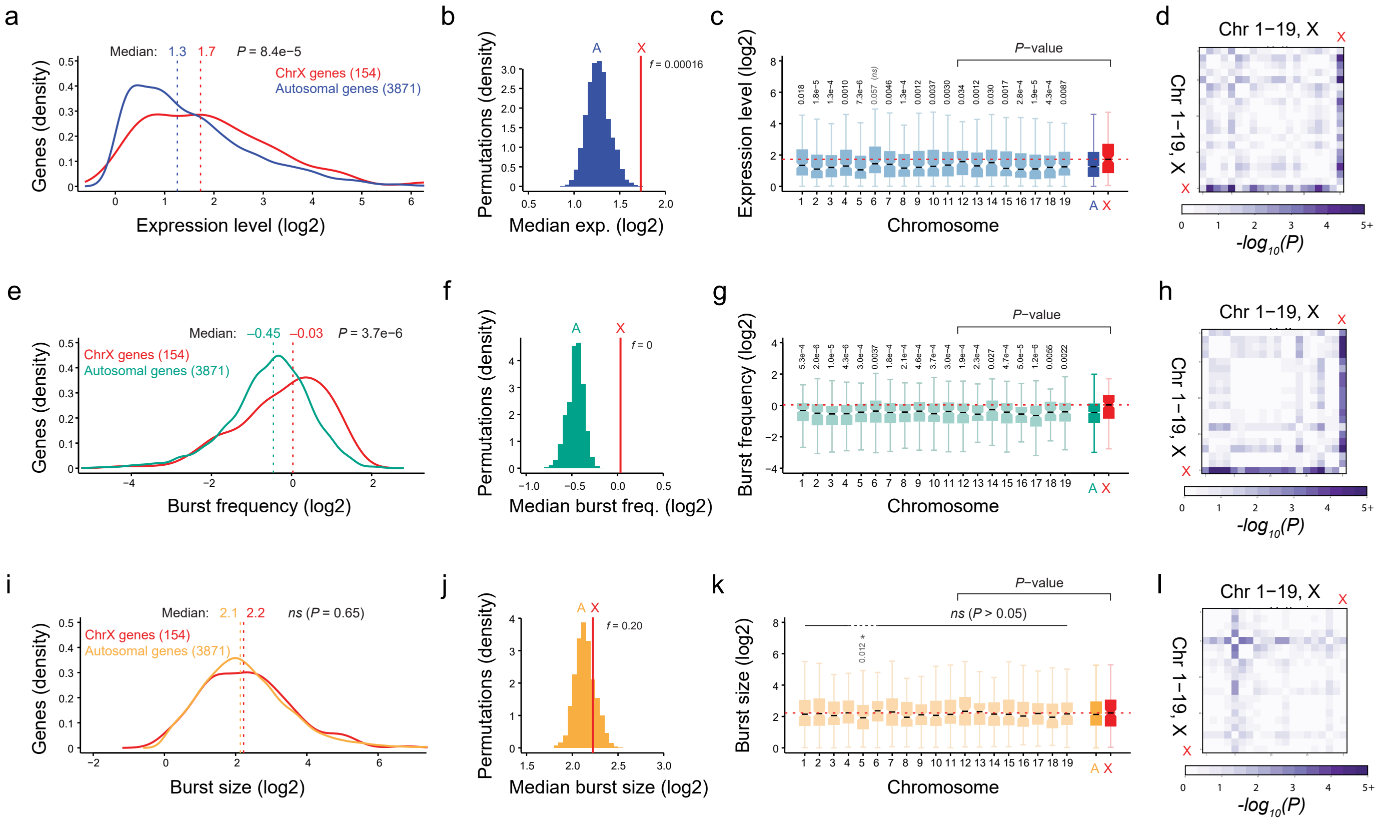


**Supplementary Figure 1. Analysis of the CAST allele in fibroblasts expressing the CAST X-chromosome.**
**(a)** Distribution of expression levels of autosomal and X-linked genes on the CAST allele in fibroblasts. Wilcoxon rank-sum test was used for significance testing. **(b)** Median expression levels of randomly selected subsets of autosomal genes compared to the X-linked genes (n=154 genes per permutation and 100,000 total permutations), and *f* denoting fraction permutations for which the autosomal median reached that of chromosome X. **(c)** Distribution of expression levels for genes on each autosomal chromosome (light blue), all autosomes (dark blue), and chromosome X (red). Centre lines denote the median; hinges denote the first and third quartiles; whiskers denote 1.5×IQR. One-sided Wilcoxon rank-sum test was used for significance testing. **(d)** *P*-values from pairwise comparisons of expression level between chromosomes from the Wilcoxon rank sum test. **(e-h)** Same as (a-d) but for burst frequency ($k_{on}$). *(i-l)* Same as (a-d) but for burst size ($k_{syn}/k_{off}$).

Supplementary figure 2

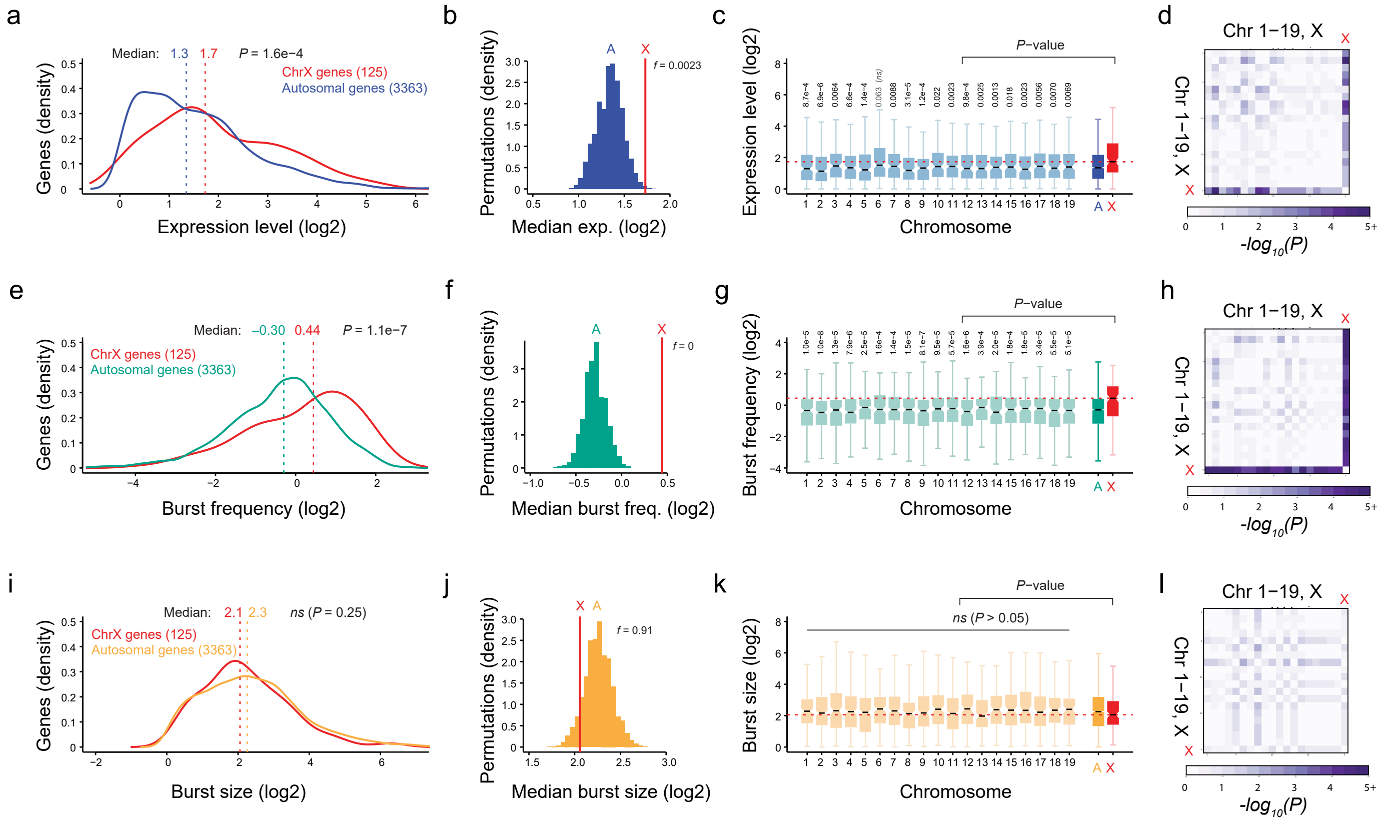


**Supplementary Figure 2. Analysis of the C57 allele in male ESCs.**
**(a)** Distribution of expression levels of autosomal and X-linked genes on the C57 allele in ESCs. Wilcoxon rank-sum test was used for significance testing. **(b)** Median expression levels of randomly selected subsets of autosomal genes compared to the X-linked genes (n=125 genes per permutation and 100,000 total permutations), and *f* denoting fraction permutations for which the autosomal median reached that of chromosome X. **(c)** Distribution of expression levels for genes on each autosomal chromosome (light blue), all autosomes (dark blue), and chromosome X (red). Centre lines denote the median; hinges denote the first and third quartiles; whiskers denote 1.5×IQR. One-sided Wilcoxon rank-sum test was used for significance testing. **(d)** *P*-values from pairwise comparisons of expression level between chromosomes from the Wilcoxon rank sum test. **(e-h)** Same as (a-d) but for burst frequency ($k_{on}$). **(i-l)** Same as (a-d) but for burst size ($k_{syn}/k_{off}$).

Supplementary figure 3

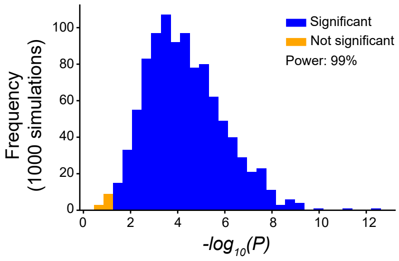


**Supplementary Figure 3. Power analysis of burst-size increase for X-linked genes.**
Distribution of significance values from the Wilcoxon rank sum test comparing the burst size ($k_{syn}/k_{off}$) of autosomal genes on the C57 allele in fibroblasts to a simulated chromosome consisting of 149 randomly selected autosomal genes which have been upregulated in expression levels to 1.4x its regular expression level by increasing burst size (n=1,000 simulations).

Supplementary figure 4

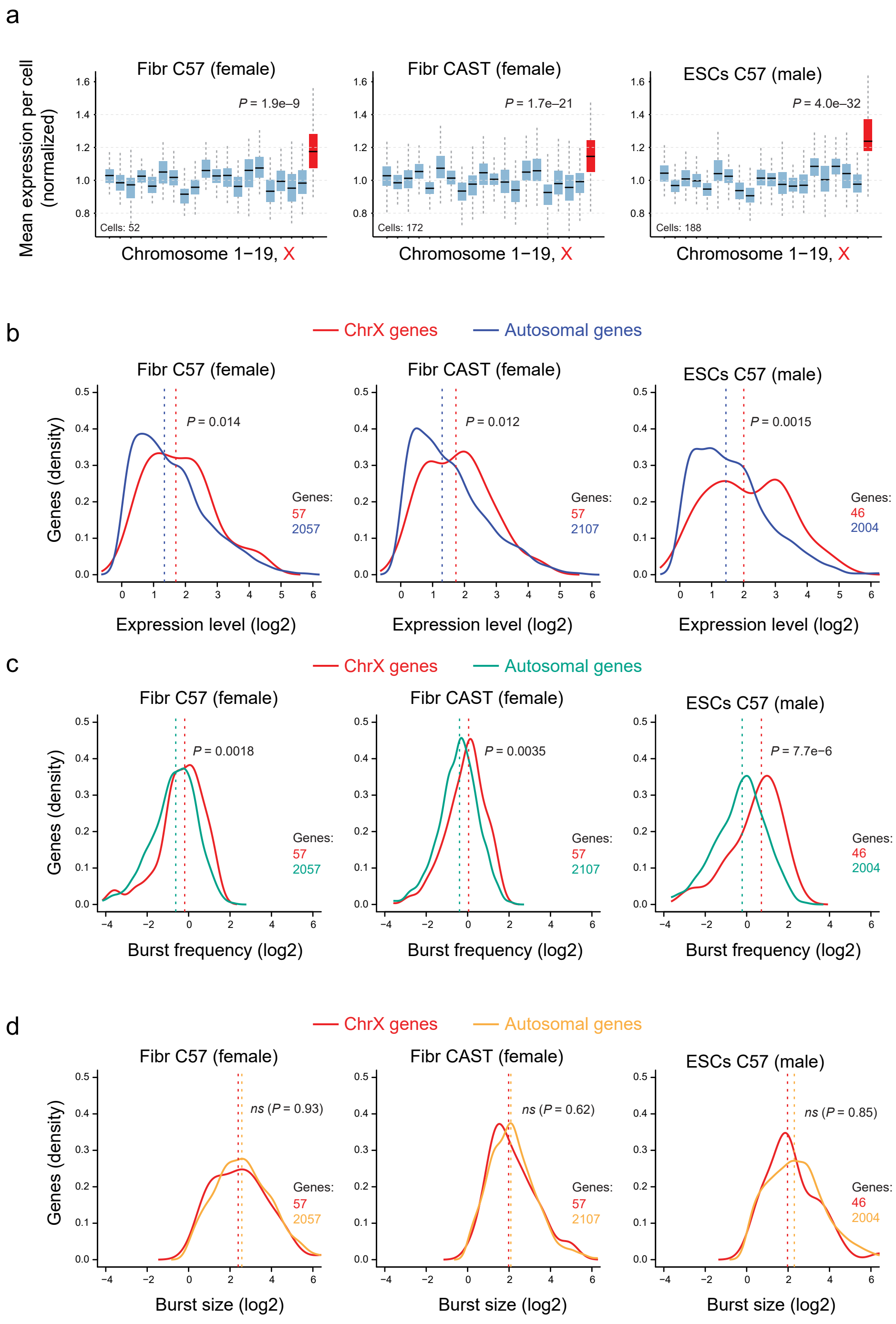


**Supplementary Figure 4. Kinetics of X-upregulation in ubiquitously expressed genes.**
**(a)** Mean expression of genes on each chromosome normalized by the mean expression of all autosomal genes in each cell and allele, restricted to ubiquitously expressed genes. Centre lines denote the median; hinges denote the first and third quartiles; whiskers denote 1.5×IQR. Wilcoxon rank-sum test was used for significance testing. **(b-d)** Distribution of (b) expression level, (c) burst frequency, and (d) burst size of ubiquitously expressed autosomal and X-linked genes for each allele and cell type. Wilcoxon rank-sum test used for significance testing.
